# Single-cell foundation models identify shared and divergent transcriptomic signatures of aging across invertebrates and mammals

**DOI:** 10.64898/2026.07.24.740647

**Authors:** Mohammad Aman Ullah Al Amin, Khoi Le, Hong Qin

**Affiliations:** Department of Computer Science, Old Dominion University, Norfolk, 23529, VA, U.S.A; Department of Computer Science, Brown University, Providence, 02912, RI, U.S.A; School of Data Science, Old Dominion University, Norfolk, 23529, VA, U.S.A

**Keywords:** single-cell RNA sequencing, aging, foundation models, SHAP, ribosomal proteins, cross-species analysis, scGPT, Geneformer, gene attribution, interpretability

## Abstract

Cross-species aging clocks have so far stayed within mammals. We asked whether a single model can predict age from single cells of fly, worm, mouse, and human, species separated by roughly 800 million years of evolution. We fine-tuned scGPT and Geneformer on 1.3 million such cells, using one parameter set per architecture. Both models classified cells as young, middle, or old, reaching 78.6% and 80.5% accuracy. SHAP attribution then showed where the two architectures agreed and where they diverged. scGPT ranked ribosomal proteins highest in every species, with RPL12 first throughout. Geneformer did the same in mouse and human but favored signaling, chromatin, and ubiquitin-ligase genes in fly and worm. Hiding all 51 ribosomal-protein genes at inference cost 22–28 percentage points of accuracy, against 0.3 points for size-matched random gene sets. Bootstrap resampling, five-fold cross-validation, and comparison with six published aging studies left the rankings largely unchanged. Age-associated signal is therefore accessible to a pooled cross-species model, but which genes carry it depends on how the model encodes expression.

## 1 Introduction

Aging involves molecular processes conserved across evolution [1, 2], yet which genes mark cellular aging in distantly related animals is still poorly resolved. Bulk transcriptomic studies point to shared changes in ribosomal, mitochondrial, and immune pathways [3, 4], and multispecies comparisons have begun to separate mechanisms of aging from mechanisms of longevity [5]. Single-cell RNA sequencing (scRNA-seq) now allows the same question at cell-type resolution, drawing on aging atlases for mouse [6], fly [7], worm [8], and human [9] that together cover millions of age-annotated cells from invertebrates and mammals. Most cross-species analyses of these data still test one gene at a time, which leaves nonlinear and context-dependent relationships among genes out of reach. Work on yeast protein-interaction networks makes a similar point from another direction: interaction structure carries evolutionary history, and age-dependent change can be described with stochastic gene networks or interaction landscapes [10–12].

Molecular clocks make a related case that chronological age is recoverable from high-dimensional omics data. The first such clocks used DNA methylation across human tissues [13] or bulk blood and fibroblast transcriptomes [14, 15]; later versions span mammalian species and tissues using methylation [16] or transcriptomes [17]. At single-cell resolution, transcriptomic clocks have quantified cell-type-specific aging and rejuvenation in mouse brain [18], and PBMC models tie predicted age to a balance between ribosome and inflammation programs [19]. All of this work stays within mammals. A clock trained on invertebrate and mammalian cells at the same time has not been reported.

Translational regulation and ribosomal protein (RP) biology have been tied to aging repeatedly and in several settings. Reduced TOR signaling extends lifespan in yeast [20], *Drosophila* [21], and mammals [22]. Depleting 60S ribosomal subunits extends yeast lifespan [23], and RNAi knockdown of RP genes does the same in *C. elegans* [24, 25]. Longitudinal transcriptomic and proteomic work has implicated protein-biogenesis machinery as an early driver of replicative aging in yeast [26], and RPL12 has been identified as a ribophagy receptor linking ribosomal function to cellular quality control [27]. Each of these results came from testing translation directly. Whether RP genes surface on their own, in an unbiased single-cell comparison of distantly related species, is a different question.

Foundation models pretrained on large scRNA-seq datasets transfer learned transcriptomic representations to new tasks [28], and cross-species pretraining can fold in shared regulatory knowledge [29]. scGPT encodes binned expression magnitudes with a generative transformer [30]; Geneformer instead orders gene tokens by expression rank [31, 32]. Because the two architectures see the same transcriptome through different representations, they amount to independent tests of any gene-level claim. Age-focused deep learning has already paired single-cell clocks with explainable AI to expose an age-associated ribosomal subnetwork in more than one million human immune cells [33]. Whether general-purpose foundation models recover comparable genes across distantly related species is open. SHAP provides one way to attribute a prediction to individual genes [34].

Here we trained scGPT and Geneformer to classify age in approximately 1.3 million cells from *Drosophila melanogaster, Caenorhabditis elegans, Mus musculus*, and *Homo sapiens*, four species separated by roughly 800 million years of evolution. For each architecture, one pooled model handled young, middle, and old cells from all four. We used SHAP to identify the genes behind those predictions, then tested the rankings by bootstrap resampling, five-fold cross-validation, comparison with six independent aging studies, and inference-time gene perturbation. scGPT returned a conserved RP signature. Geneformer, working from the same cells, returned signaling and proteostasis genes instead, mostly in the invertebrates.

## 2 Results

### 2.1 Foundation models achieve accurate cross-species age classification

We combined single-cell RNA-seq aging atlases from fly, worm, mouse, and human (Table 1, Fig. 1). Species-specific ages were assigned to young, middle, or old categories, and non-human genes were mapped to one-to-one human orthologs with DIOPT. This left approximately 1.3 million cells and a shared set of 2,337 genes.

**Table 1.**
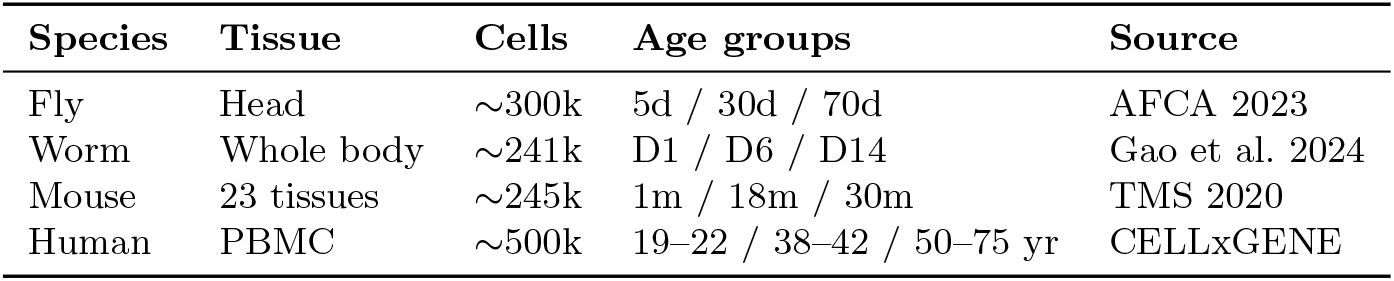
Summary of multispecies aging datasets.

| Species | Tissue | Cells | Age groups | Source |
| --- | --- | --- | --- | --- |
| Fly | Head | ~300k | 5d / 30d / 70d | AFCA 2023 |
| Worm | Whole body | ~241k | D1 / D6 / D14 | Gao et al. 2024 |
| Mouse | 23 tissues | ~245k | 1m / 18m / 30m | TMS 2020 |
| Human | PBMC | ~500k | 19–22 / 38–42 / 50–75 yr | CELLxGENE |

**Fig. 1.**
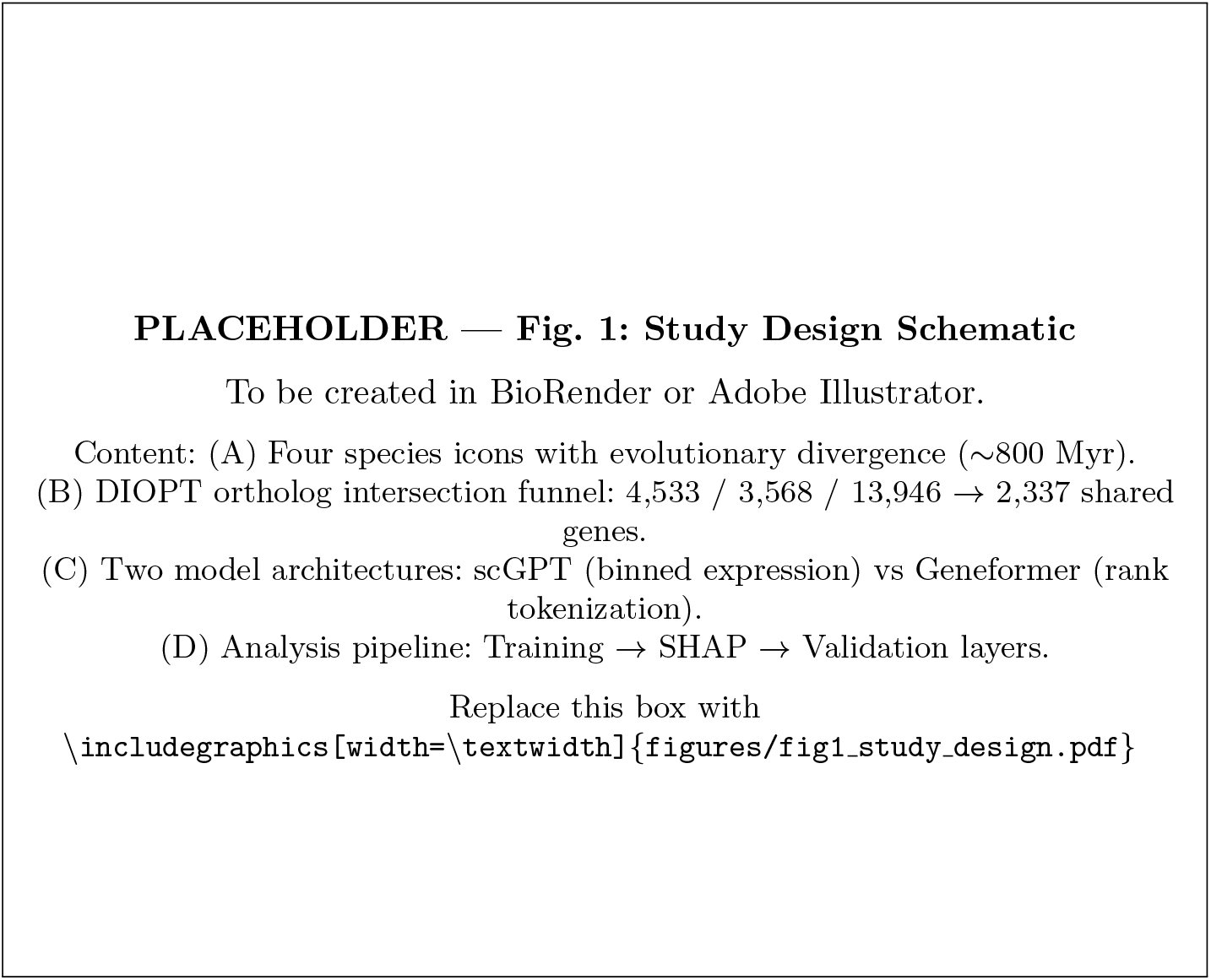
Study design and analysis pipeline. **(A)** Single-cell RNA-seq aging atlases from four species span approximately 800 million years of evolution. **(B)** DIOPT one-to-one ortholog mapping defines a shared set of 2,337 genes. **(C)** We fine-tuned scGPT and Geneformer to classify cells as young, middle, or old. **(D)** We used SHAP to rank genes and tested the rankings by resampling, cross-validation, literature comparison, and inference-time gene perturbation.

Fine-tuning used a non-donor-aware (NDA) protocol, in which cells rather than donors were split between training and test sets (Table 2). A four-stage search over 66 scGPT configurations showed that learning rate (1 *×* 10^*−*5^) and batch size (16) mattered most (Supplementary Fig. S1); Geneformer ran with its default fine-tuning configuration. Per-species accuracy ranged from 77.9% in human to 96.0% in mouse for scGPT, and from 88.1% to 94.7% over the same two species for Geneformer (Fig. 2). Human was the hardest case for both models, which is unsurprising given 25 young donors against 147 old ones and a continuous spread of adult ages. A single pooled model reached 78.6% (scGPT) and 80.5% (Geneformer) across all four species, using one parameter set and a shared vocabulary of 2,337 genes. Holding out complete donors was considerably harder and dropped accuracy to 51–73% (Supplementary Table S1); training on one species and testing on another left both models near chance (29–41%).

**Table 2.** NDA test accuracy and macro F1 for scGPT and Geneformer across four species and the Combined (pooled) model.

| Species | scGPT Acc (%) | scGPT F1 | GF Acc (%) | GF F1 |
| --- | --- | --- | --- | --- |
| Fly | 93.4 | 0.926 | 94.6 | 0.939 |
| Worm | 87.7 | 0.867 | 91.0 | 0.889 |
| Mouse | 96.0 | 0.958 | 94.7 | 0.945 |
| Human | 77.9 | 0.775 | 88.1 | 0.880 |
| Combined | 78.6 | 0.782 | 80.5 | 0.793 |

**Fig. 2.**
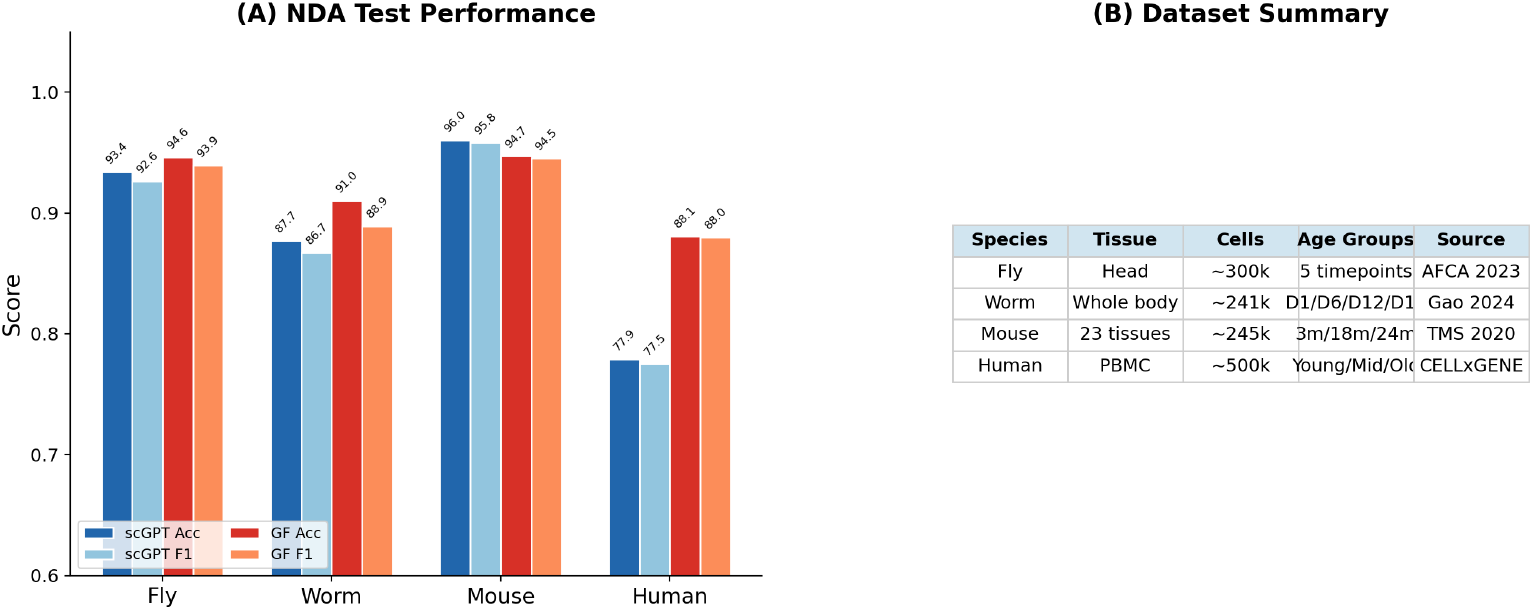
Model performance across species. Test accuracy and normalized confusion matrices for scGPT and Geneformer under the NDA protocol. Most errors fall between adjacent age categories rather than between young and old.

### 2.2 scGPT identifies a conserved ribosomal protein aging signature

We applied SHAP GradientExplainer [34] to the fine-tuned, pooled NDA classifiers to identify genes that contributed to age predictions. RP genes dominated the scGPT rankings, accounting for 70–80% of the top 10 genes and 46–52% of the top 50 in each species (Fig. 3). RPL12 ranked first in all four species, and the same four genes (RPL12, RPS21, RPS13, and RPS26) held ranks 1–4 everywhere.

**Fig. 3.**
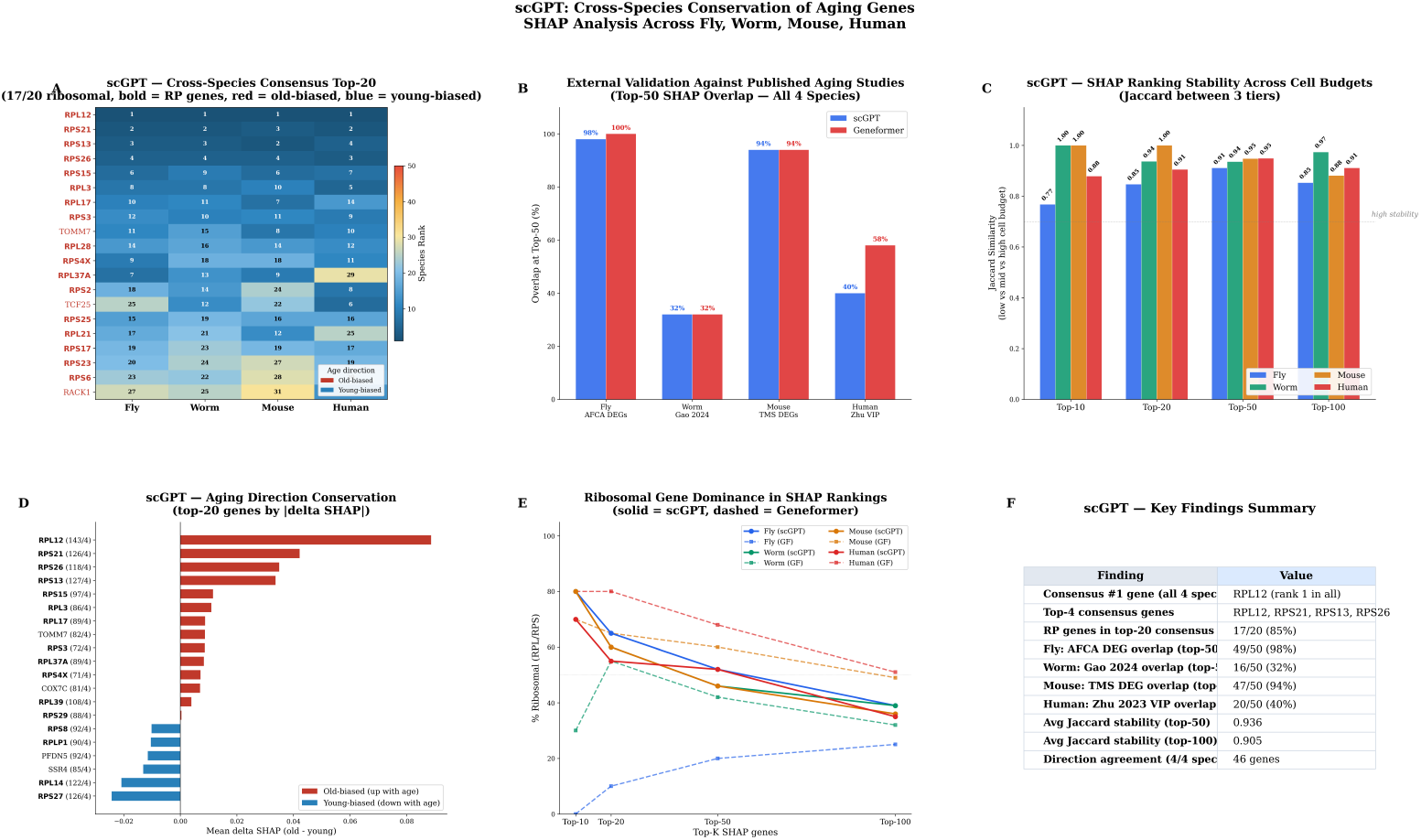
scGPT SHAP gene rankings across four species. The plot shows the top-ranked genes from the pooled NDA model and marks RP genes by color. RPL12 ranks first in all four species; RPL12, RPS21, RPS13, and RPS26 occupy the top four positions in each species.

Twenty-four genes appeared in the top 50 of all four species at once (Fig. 4), far more than chance would give. Most were ribosomal: RPL12, RPS21, RPS13, RPS26, RPS15, and RPL3 among them. The conserved non-ribosomal genes were TOMM7, TCF25, HECTD1, and RB1CC1, which act in mitochondrial transport, nutrient sensing, ubiquitination, and autophagy. Old-biased genes were more conserved across species (Jaccard 0.52–0.56) than young-biased genes (0.32–0.47), although a ranking comparison of this kind says nothing about when the underlying programs arose.

**Fig. 4.**
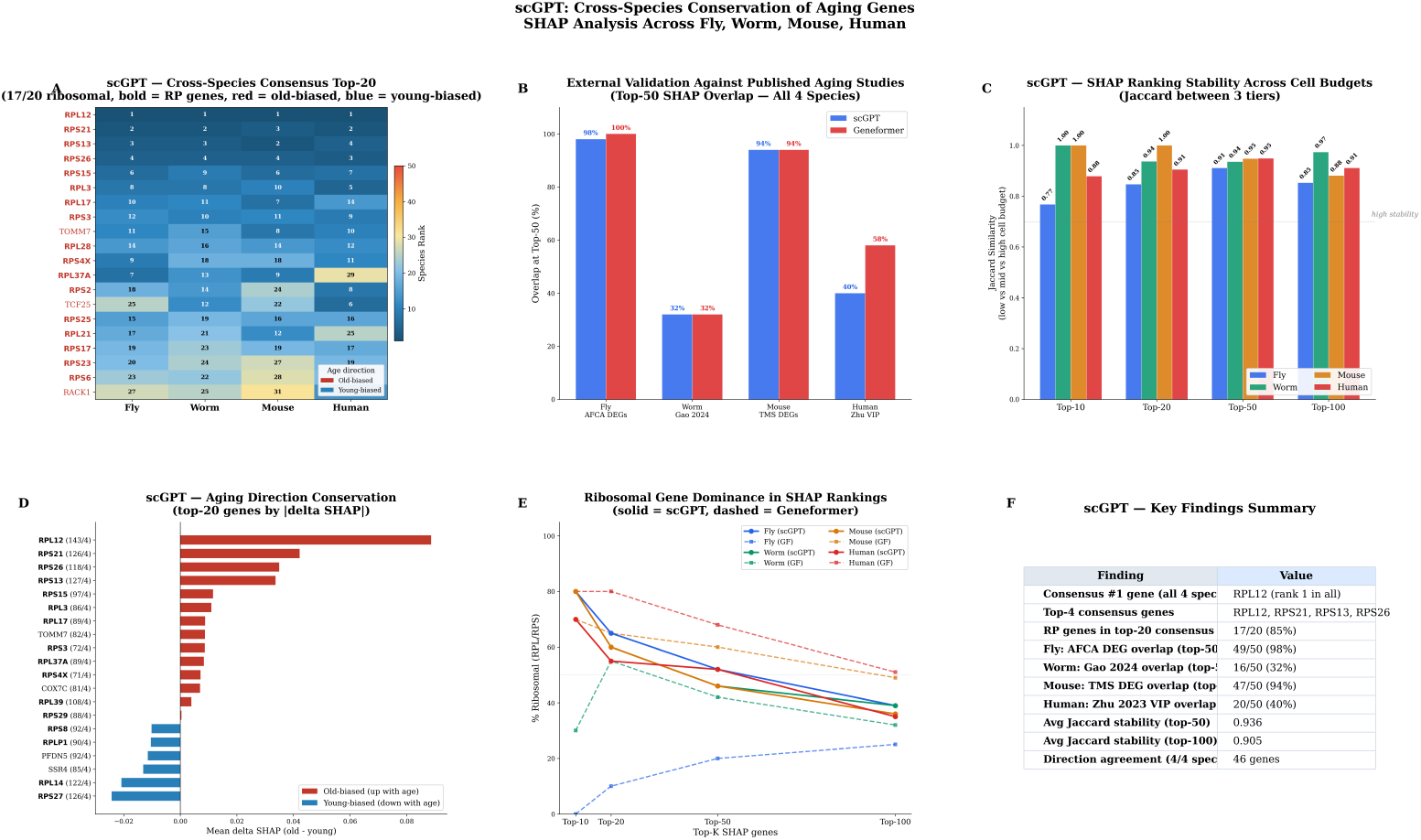
Cross-species conservation of scGPT SHAP rankings. Twenty-four genes appear in the top 50 of all four species simultaneously. Old-biased genes show higher cross-species Jaccard similarity than young-biased genes.

### 2.3 Geneformer prioritizes distinct aging biology in invertebrates

In the two invertebrates, Geneformer produced a different picture (Fig. 5). Its five highest-ranked fly genes (PIP4K2A, ELF2, PLCB4, TCF7L2, and NIPBL) act in phosphoinositide signaling, transcriptional regulation, and chromatin organization, and no RP gene reached the fly top 10. In worm, three of the top five (HECTD1, UBR5, and HUWE1) encode E3 ubiquitin ligases, although RP genes still appeared among the worm top 20, so the shift away from translation was a gradient from mammals to invertebrates rather than an absolute absence.

**Fig. 5.**
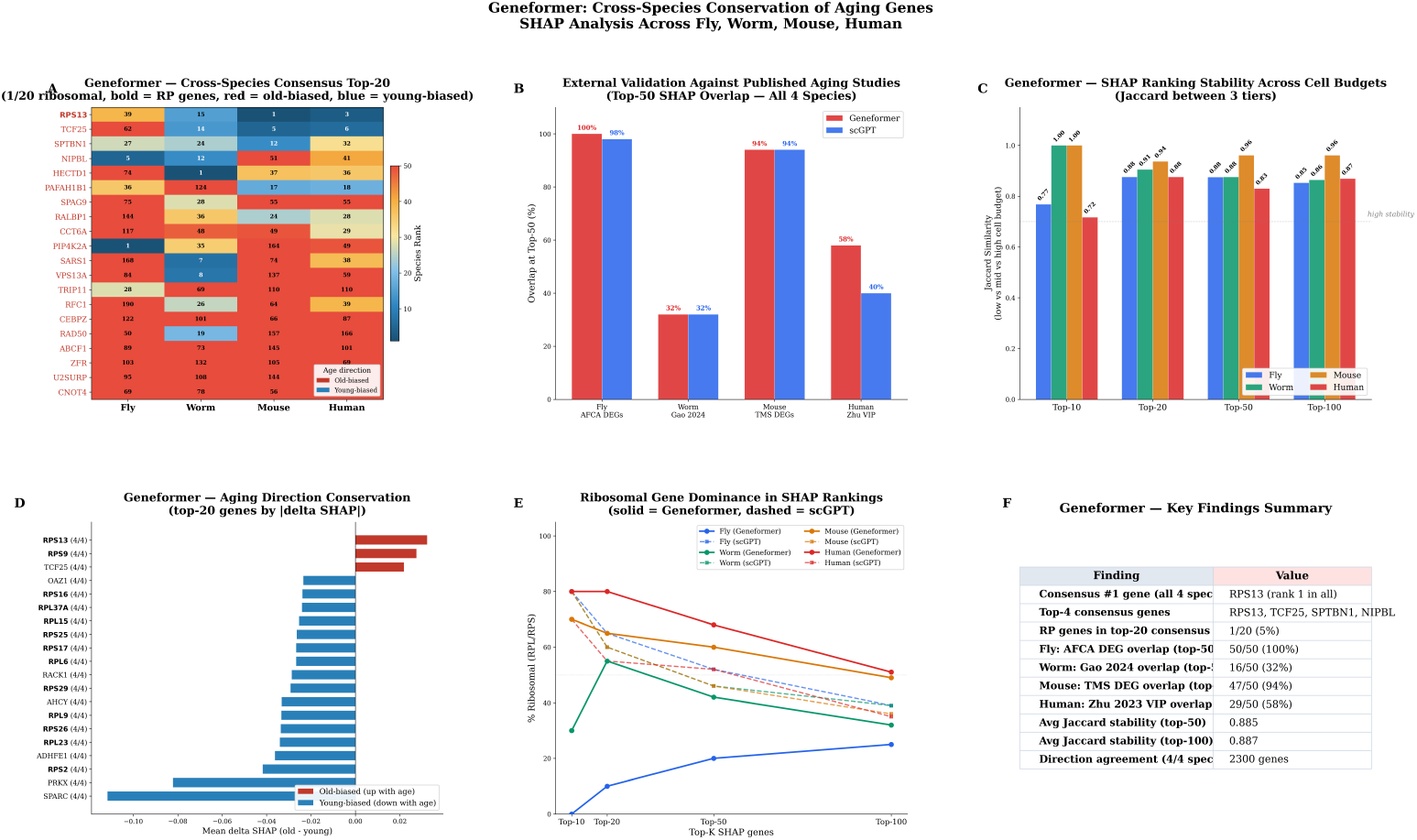
Geneformer SHAP gene rankings across four species. Signaling, chromatin, and ubiquitin-ligase genes lead the fly rankings, with no RP genes among the fly top 10; worm rankings retain some RP genes but are dominated by E3 ubiquitin ligases. RP genes lead the mammalian rankings. The shift in lineage-dependent importance among conserved orthologs runs as a gradient from mammals to invertebrates. RPS13 ranks first in the cross-species consensus.

The mammalian rankings were another matter, and closely resembled scGPT’s. Three of the five highest-ranked mouse genes (RPS13, RPS9, and RPS29) and four of the five highest-ranked human genes (RPS9, RPS20, RPS13, and RPS27) encode ribosomal proteins. The direction of the effect split the same way: Geneformer called 48 of 51 RP genes young-biased in the invertebrates, whereas scGPT’s directions varied by species. Pooled across species, Geneformer ranked RPS13 first and TCF25 second; TCF25 has been described as a nutrient sensor acting on V-ATPase activity and autophagy [35].

### 2.4 Cross-model comparison identifies shared and architecture-dependent signals

The two sets of rankings agreed only in part. Top-50 lists shared 9–17 genes per species, with rank correlations of *ρ* = 0.21 to 0.51 (Fig. 6). RPS13 was the only gene to reach both models’ top 50 in all four species. Some of the disagreement probably traces to the input representations, since scGPT keeps binned expression magnitude while Geneformer keeps only the rank order of expressed genes.

**Fig. 6.**
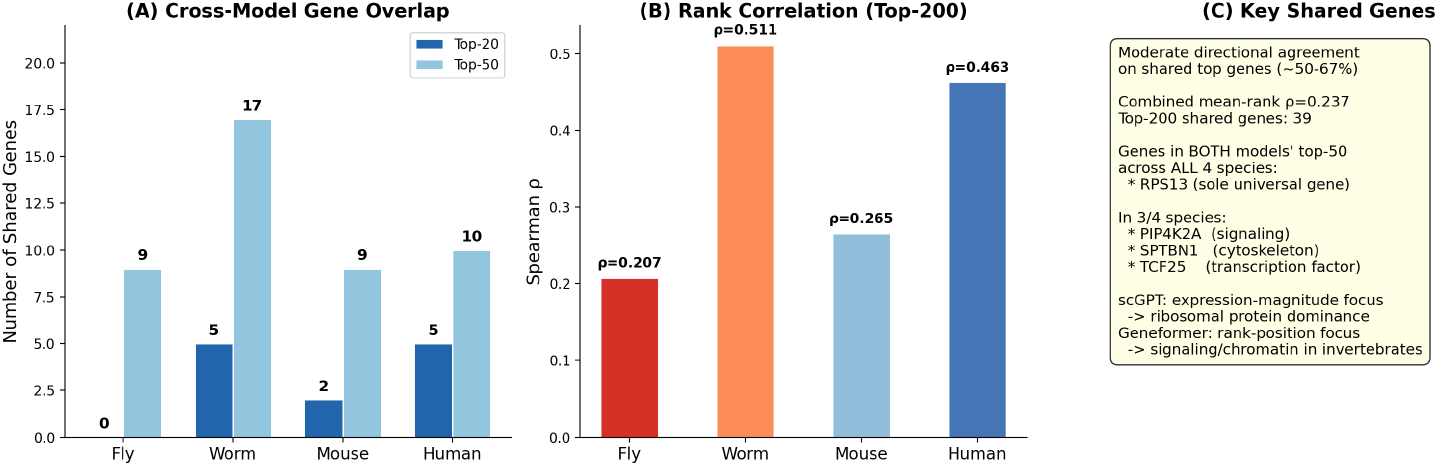
Cross-model agreement between scGPT and Geneformer. Top-50 overlap ranges from 9 to 17 genes per species. RPS13 is the sole gene in both models’ top 50 across all four species.

Pathway-level agreement was better than gene-level agreement. Gene Ontology (GO) analysis at FDR *<* 0.05 recovered 6 of the 12 Lopez-Otín hallmarks of aging across models and species (Fig. 7), with loss of proteostasis the strongest and cytoplasmic translation reaching *p <* 10^*−*9^. Three-class SHAP heatmaps showed how the signals shifted with age (Fig. 8): in scGPT, mammalian RP genes usually reached their largest values at middle age, while Geneformer tied RP genes more consistently to youth. Both models again separated invertebrates from mammals by direction.

**Fig. 7.**
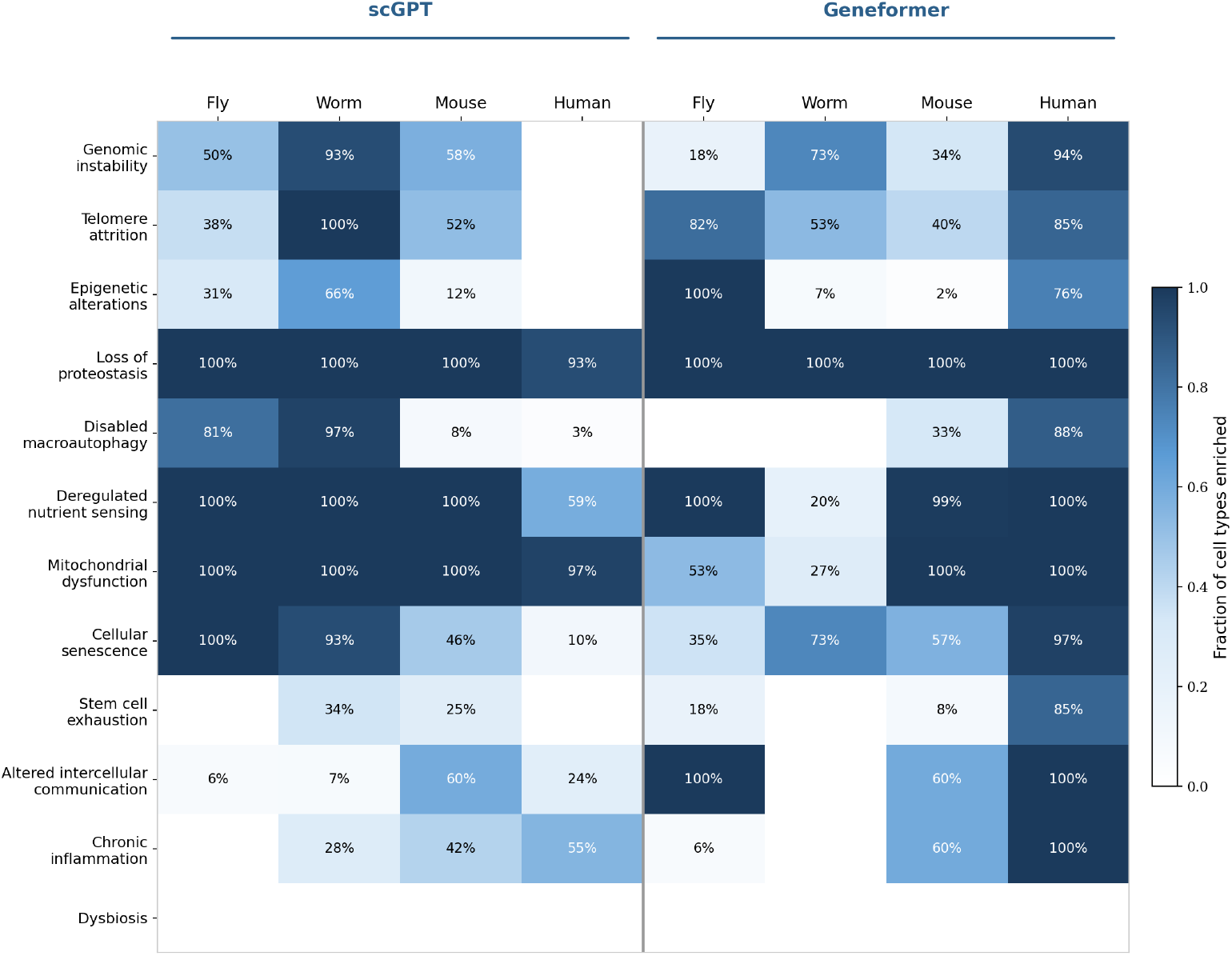
GO biological process enrichment mapped to the 12 hallmarks of aging. Enriched terms map to 6 of the 12 hallmarks across the models and species. Loss of proteostasis has the strongest enrichment.

**Fig. 8.**
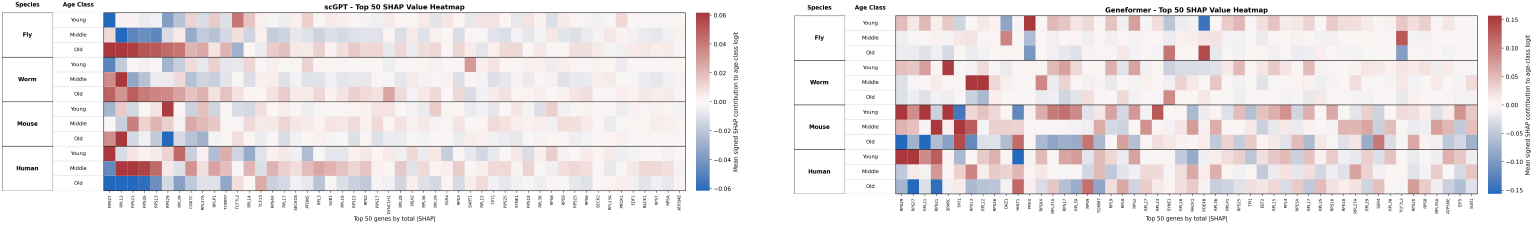
Signed SHAP heatmaps across the three-class aging trajectory. **Left:** scGPT. **Right:** Geneformer. Mammalian RP genes in scGPT often peak in middle age, while Geneformer shows RP genes more uniformly associated with youth.

### 2.5 Independent analyses support ranking robustness

We checked the rankings four ways (Fig. 9, Fig. 10).

**Fig. 9.**
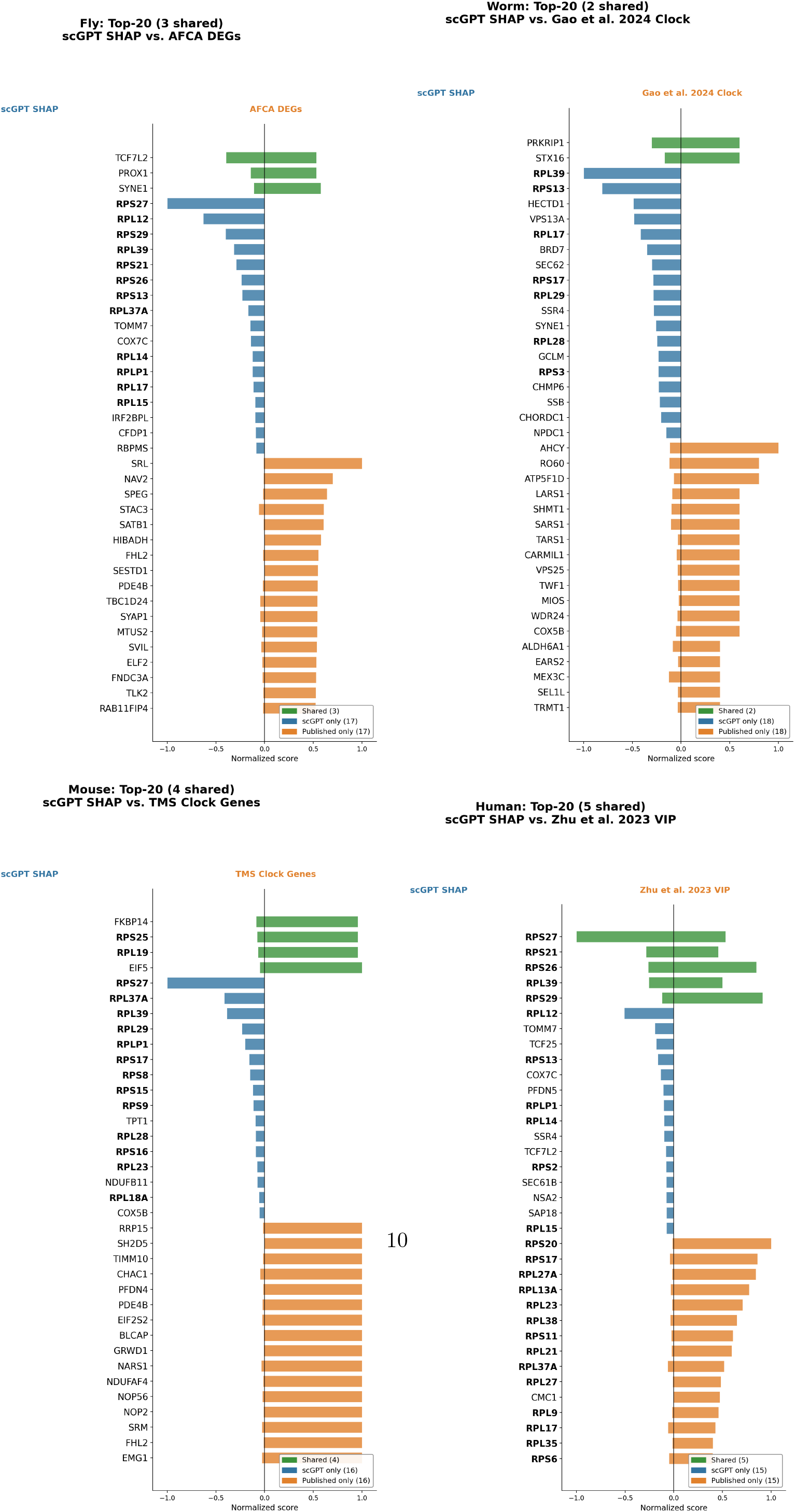
Comparison of SHAP gene rankings with independent aging studies. The plots show overlap between the top 50 SHAP genes and published aging gene sets for each species. Human has the strongest enrichment (10–15-fold) but the fewest overlapping genes.

**Fig. 10.**
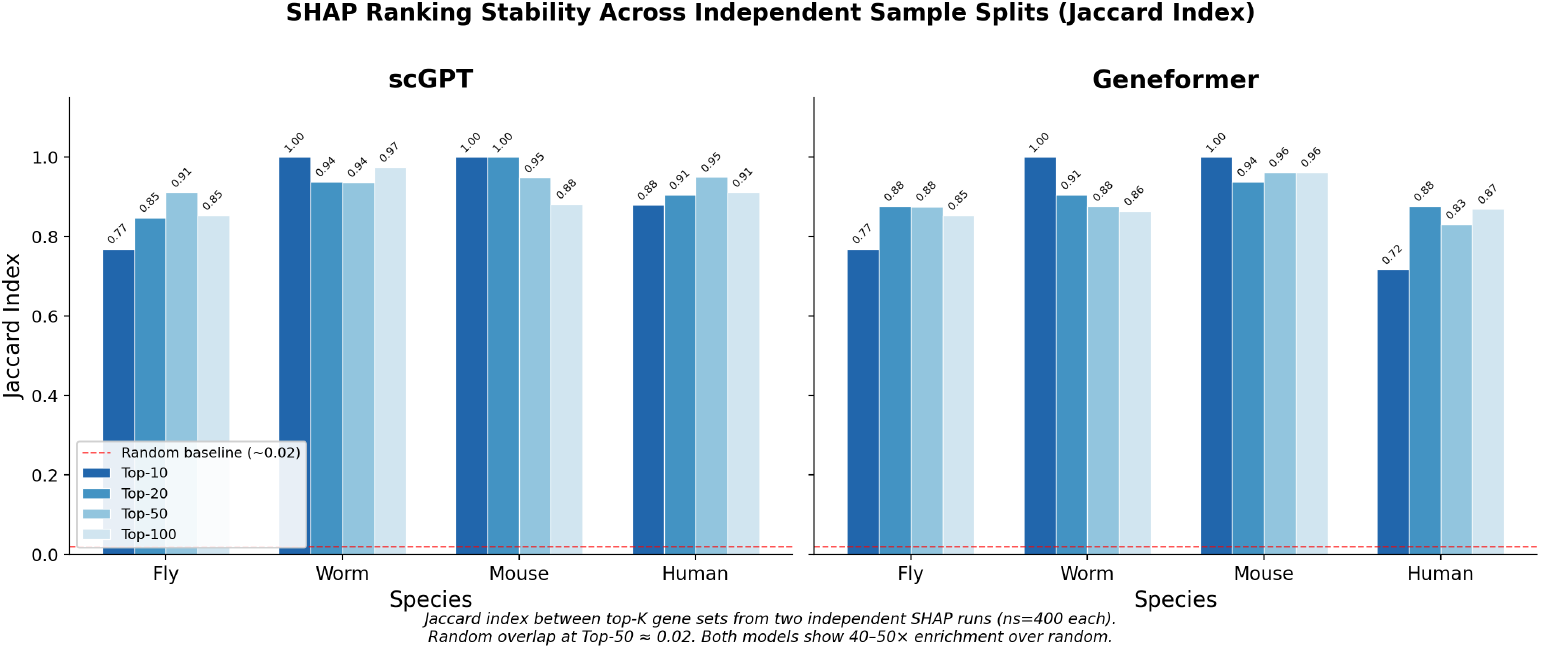
SHAP ranking stability. Both models have a median Jaccard index above 0.87, more than 40-fold above the random baseline.

#### Bootstrap resampling

Across 1,000 resamples of the explained cells, RPL12 held first place in every scGPT resample (SD = 0.00; 95% CI [1, 1]), and Geneformer’s top gene, RPS13, moved by an SD of 0.13 ranks. Combining bootstrap stability with cross-species conservation, cross-model agreement, and literature support gave 33 high-confidence candidates, 18 of them Tier 1 and 15 Tier 2.

#### Cross-validation

Retraining each model in every fold of a five-fold cross-validation left ranking stability 26-fold (scGPT) and 43-fold (Geneformer) above the permutation null (Table 3). Geneformer scored higher on the Nogueira index (0.63 versus 0.43) and on directional consistency (0.99 versus 0.82). Seven genes were stable in both models and kept the same age direction: RPS13, RPS21, RPS26, RPS27, RPS8, SPARC, and TCF25.

**Table 3.** Cross-validation stability metrics at *K* = 50 (harmonized absolute-delta ranking).

| Metric | scGPT | Geneformer |
| --- | --- | --- |
| Nogueira stability | 0.43 | 0.63 |
| Pairwise Jaccard | $0.29 \pm 0.04$ | $0.47 \pm 0.07$ |
| Observed/Null ratio | 26× | 43× |
| CV-stable genes | 20 | 33 |
| Direction consistency | 0.82 mean | 0.99 mean |

#### Literature comparison

Against gene sets from six independent aging studies, the SHAP rankings were enriched 2–15-fold. The fly rankings covered 98–100% of AFCA differentially expressed genes (DEGs) and the mouse rankings 94% of Tabula Muris Senis DEGs. Human overlap was smaller in absolute terms but enriched 10–15-fold against the PBMC aging genes reported by Zhu et al., whose list is short to begin with. Every top-10 gene had a prior link to aging, and 53 of 60 hypergeometric tests reached *p <* 0.05.

#### Independent cell subsets

Recomputing SHAP values on non-overlapping subsets of cells gave Jaccard indices of 0.77 to 1.00 for scGPT and 0.72 to 1.00 for Geneformer, more than 40-fold above the random baseline.

### 2.6 Inference-time gene perturbation shows model reliance on RP genes

To ask whether the trained models actually relied on RP genes, we hid selected genes from the test input and left the model weights untouched (Table 4; Fig. 11). For scGPT this meant zeroing the target genes before tokenization; for Geneformer, dropping their tokens from the rank-ordered sequence.

**Table 4.** Inference-time gene perturbation results. Random controls report mean *±* SD across 5 gene sets.

| Condition (genes hidden) | scGPT | $\Delta$ | GF | $\Delta$ |
| --- | --- | --- | --- | --- |
| Baseline (0) | 81.7% | — | 80.5% | — |
| RP top 50 (25) | 74.1% | −7.7 pp | 74.4% | −6.1 pp |
| All RP (51) | 54.1% | −27.6 pp | 58.4% | −22.1 pp |
| Top 50 (50) | 73.4% | −8.4 pp | 74.4% | −6.1 pp |
| Top 100 (100) | 73.7% | −8.0 pp | 74.4% | −6.1 pp |
| Random 51 (51) | 81.4% | −0.3 pp | 80.2% | −0.3 pp |

**Fig. 11.**
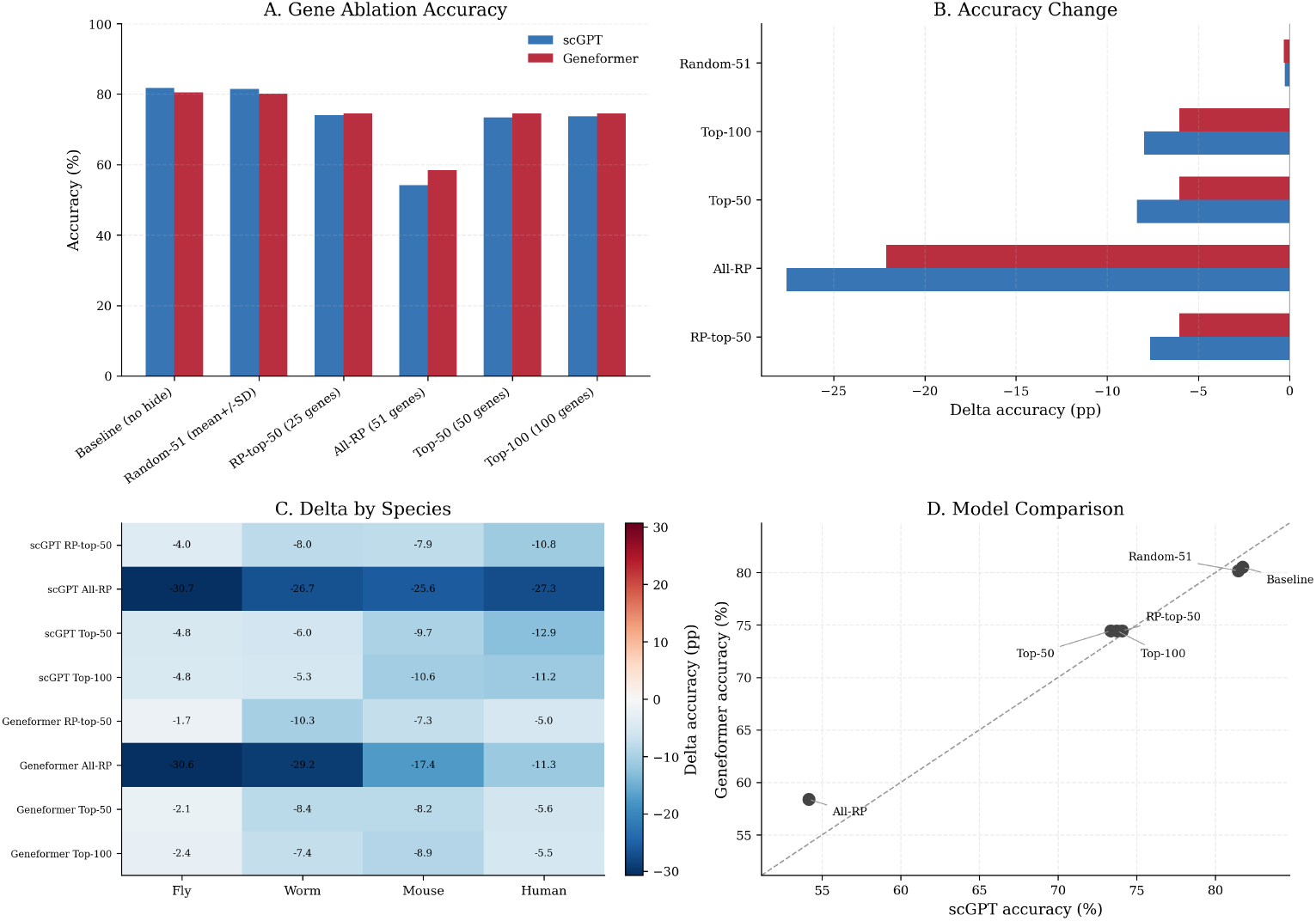
Inference-time gene perturbation. Hiding all 51 RP genes reduces accuracy by 22–28 percentage points, compared with 0.3 points for random controls (*Z >* 25). Removing the 51 RP genes has a larger effect than removing the top 100 SHAP genes.

Hiding all 51 RP genes cost scGPT 27.6 percentage points (81.7% to 54.1%) and Geneformer 22.1 points (80.5% to 58.4%). Size-matched random controls cost 0.3 points (*Z >* 25, *p ≪* 0.001). The 51 RP genes also cost more than the top 100 SHAP genes, which took 8.0 points off scGPT and 6.1 off Geneformer. Whatever the models had learned, it leaned on the RP set as a group more than on a larger collection of individually high-ranking genes.

Retraining after removing the RP genes told a different story (Supplementary Section S2): accuracy fell by only 3.1 and 5.4 points, so both models could partly compensate when given the chance to relearn. The inference-time losses were fourfold to ninefold larger.

## 3 Discussion

Both architectures classified relative life stage across four species from a single parameter set, which was not a given: these cells share only 2,337 mappable genes and nothing resembling a common aging timescale. What the two models attended to, however, was not the same. scGPT put RPL12 first in every species and RP genes throughout its rankings, a result that sits comfortably with decades of work connecting reduced translation to longevity, from TOR signaling in yeast and *Drosophila* [20, 21] to RP gene knockdown in *C. elegans* [24, 25] and RPL12-mediated ribophagy [27]. Yeast proteomics shows coordinated age-related change in protein biosynthesis, stress responses, and mitochondrial proteins [36], and mammalian transcriptomes carry conserved ribosomal and translational changes [17]. Geneformer, given the same cells, prioritized RP genes only in the mammals and turned to signaling, chromatin, and ubiquitin-ligase genes in the invertebrates. We read the RP signal, then, as an age-associated program whose weight in a prediction depends on the species and on how expression is encoded.

That dependence on encoding is worth taking seriously, because it is also what makes agreement between the models informative. scGPT discretizes expression into per-cell quantile bins, preserving ordinal magnitude within each cell; Geneformer normalizes by a fixed human-corpus median and keeps only the resulting rank order. A gene that both models place near the top is therefore unlikely to be an artifact of either representation. RPS13 was the only such gene in all four species, and TCF25 came close, ranking highly in both models in three of the four. TCF25 senses nutrients and regulates V-ATPase activity and autophagy [35], which puts it at the junction of two established features of aging. The same filters also surfaced SPAG9, a JNK scaffold; CIR1, a chromatin regulator; and VPS13A, a lipid-transport protein. Splitting age into three classes rather than two added something as well: mammalian RP genes in scGPT tended to peak at middle age, while in Geneformer they stayed youth-associated throughout, so even the apparent timing of the RP program shifts with the representation.

The rankings survived the checks we could apply. RPL12 never left first place under resampling, cross-validation stability ran 26–43-fold above the permutation null, six independent studies gave 2–15-fold enrichment, and every top-10 gene already had a documented link to aging. The perturbation experiment is the strongest of these, but its scope is narrow: it shows that the models depend on RP genes, not that RP genes cause organismal aging. We avoided cell-level differential expression as a primary check because pseudoreplication inflates its test statistics [37, 38], and leaned instead on published gene sets, cross-species conservation, resampling, retraining, and perturbation.

Several things limit what these results can support. Keeping only one-to-one DIOPT orthologs discards genes with complex evolutionary histories, where cross-species integration based on protein-language-model embeddings would retain a broader set [39]. The 2,337-gene vocabulary that remains is small enough that RP genes may double as species markers, and models trained separately on each species did show less RP dominance when their vocabularies were larger. Because the NDA protocol splits cells rather than donors, donor-and batch-associated variation can leak into the SHAP values; holding out complete donors cost 20–45 percentage points of accuracy. Tissue composition differs across the datasets, and most species are represented by a single tissue or tissue collection. The human PBMC data are unbalanced at 25 young against 147 old donors, and although the human rankings stayed stable (Jaccard 0.88–0.95) with 10–15-fold literature enrichment, stability is not a remedy for a sampling limitation. The models also predict chronological life-stage categories, and we have not tested them as biomarkers of biological age against functional outcomes, mortality, or response to longevity interventions [40]. Finally, we explained scGPT logits and Geneformer softmax outputs, so part of the divergence between the two may be a matter of target rather than architecture.

Seven genes (RPS13, RPS21, RPS26, RPS27, RPS8, SPARC, and TCF25) held their rank across both architectures and all cross-validation folds without changing age direction. They are the obvious place to start experimental work. Adding species and tissues, balancing donors, and perturbing these candidates in vivo would show how much of the model-derived signal reflects mechanism rather than the particular data and encodings used here.

## 4 Methods

### 4.1 Datasets and preprocessing

Single-cell RNA-seq aging atlases were assembled for four species (Table 1; Fig. 12). The worm data came from GEO (GSE229022), restricted to the N2D subset, with ages parsed from the D1, D6, D12, and D14 identifiers. Fly data came from the Aging Fly Cell Atlas, and mouse data from Tabula Muris Senis, for which we concatenated cells from 23 tissue-specific AnnData files. Human PBMC data came from the Asian Immune Diversity Atlas through CZ CELLxGENE Discover; we mapped Ensembl identifiers to gene symbols, summed counts where several identifiers mapped to one symbol, and kept the genes present in the scGPT vocabulary. All datasets were converted to AnnData format and labeled by species.

**Fig. 12.**
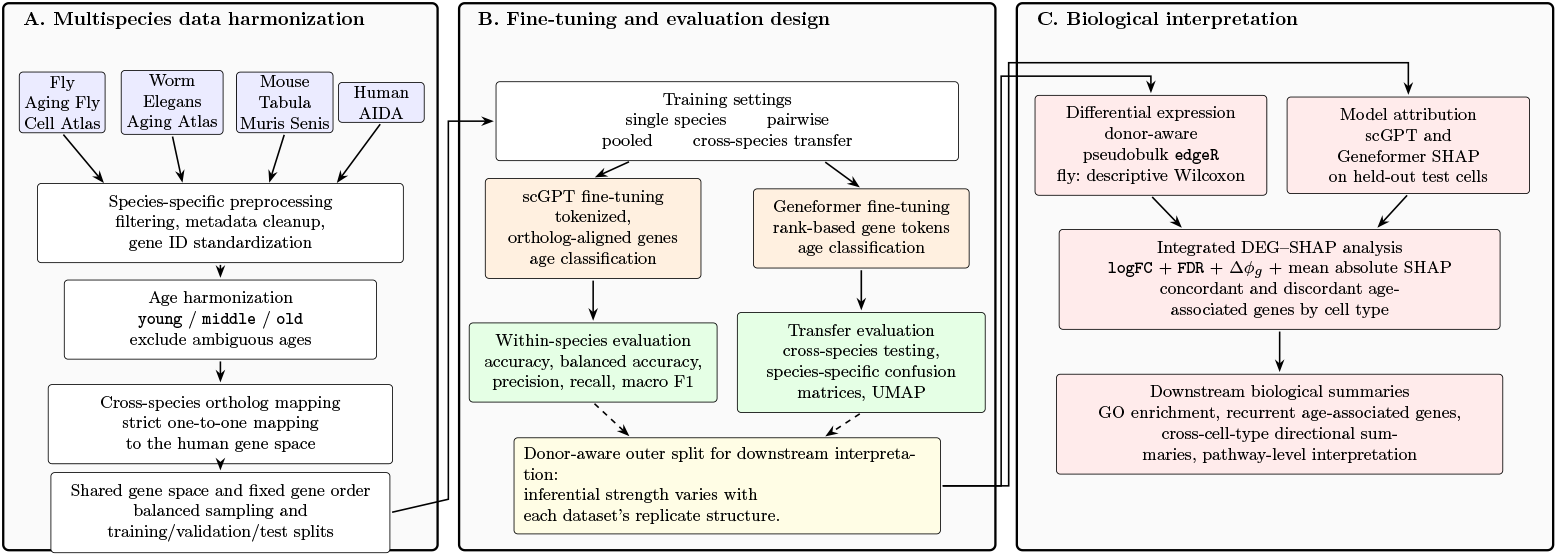
Schematic overview of the multispecies single-cell aging framework. The workflow assigns fly, worm, mouse, and human scRNA-seq data to common age classes and aligns them in a shared ortholog space. We fine-tuned scGPT and Geneformer on single species, species pairs, all four species, and cross-species transfer tasks. We combined differential expression with SHAP attribution to identify age-associated genes and pathways.

### 4.2 Cross-species gene mapping and age harmonization

Non-human genes were mapped to human orthologs using DIOPT reference mappings, keeping strict one-to-one relationships only. For each training configuration we took the shared genes and reordered every dataset so that a given input position always held the same ortholog.

Ages were collapsed into young, middle, and old on a species-specific schedule: 5, 30, and 70 days for fly; days 1, 6, and 14 for worm; 1, 18, and 30 months for mouse; and 19–22, 38–42, and 50–75 years for human. Intermediate ages, and human ages outside these ranges, were excluded.

### 4.3 Balancing and data splitting

Human cells were downsampled to equalize the three age groups, then every species was downsampled to the size of the smallest, so that pooled training was not dominated by any one of them. Splits of 81% training, 9% validation, and 10% test were drawn with fixed random seeds.

### 4.4 Model training

scGPT hyperparameters came from a four-stage, coarse-to-fine grid search of 66 runs (Supplementary Fig. S1). The final configuration used a learning rate of 1 *×* 10^*−*5^, a batch size of 16, 512-dimensional embeddings, 12 transformer layers, 8 attention heads, dropout of 0.2, a mask ratio of 0.0, no masked-language-model objective, and 30 training epochs. Expression values were discretized into 51 bins, and a `<cls>` token was prepended to each cell sequence. Geneformer used its default fine-tuning configuration and rank-based gene tokenization. Both models started from pretrained checkpoints: the whole-human checkpoint for scGPT, V2-104M for Geneformer [32].

Three training configurations were run: single species, all six species pairs, and all four species pooled. Each run kept the checkpoint with the lowest validation loss, and we report accuracy, per-class precision and recall, and macro-averaged F1 on the held-out test set. NDA models split cells across partitions; donor-aware (DA) models assigned complete donors to the test set (Supplementary Table S1).

### 4.5 SHAP analysis

SHAP values were computed for held-out test cells with GradientExplainer applied to each fine-tuned classifier. The background reference was sampled from training cells and balanced across age groups and cell types, and the explained test cells were balanced across donors or samples so that no single biological replicate dominated the estimates. We explained the old-class output, so positive SHAP values push a prediction toward old and negative values toward young.

Two summary quantities were calculated per gene:

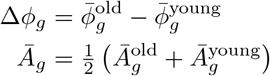

where Δ*ϕ*_*g*_ is the signed SHAP difference between old and young cells for gene *g* and *Ā*_*g*_ is the mean absolute SHAP value across cells. Cross-species comparisons ranked genes by |Δ*ϕ*_*g*|_.

Top-ranked SHAP gene sets were tested for GO enrichment with gProfiler at FDR *<* 0.05, using the corrected ortholog vocabulary (*∼*2,253 genes) as background. Enriched terms were mapped to the 12 Lopez-Otín hallmarks of aging with a predefined and deliberately conservative keyword classifier.

### 4.6 Ranking stability and bootstrap uncertainty quantification

Ranking stability was measured on two independent, non-overlapping subsets of 400 test cells per species, taking Jaccard indices between the top 10, 20, and 50 genes of each subset. Explained cells were then resampled 1,000 times within species to give each gene a mean rank, rank SD, 95% CI, and top-50 frequency. Confidence intervals for model performance used 10,000 bootstrap iterations.

### 4.7 Cross-validation stability analysis

Five-fold cross-validation used StratifiedKFold (`n_splits=5, shuffle=True, random_state=42`) with a compound species *×* age stratification key. Within each fold, both models were reinitialized from their pretrained checkpoints, retrained, and re-explained on the fold-specific test partition. Stability was scored by pairwise Jaccard index, the Nogueira stability index, and direction consistency, the fraction of folds agreeing on old versus young bias. A further 500 within-fold, species-stratified bootstrap resamples separated cell-resampling variability from variability of the training procedure as a whole.

### 4.8 High-confidence gene tiering

Genes were sorted by four criteria: bootstrap stability (top-50 frequency *≥*95% across 1,000 resamples), cross-species conservation (top 50 in at least two species), cross-model agreement (top 50 in both scGPT and Geneformer), and literature support (overlap with at least one independent aging gene set). Tier 1 genes met all four; Tier 2 met three.

### 4.9 External literature comparison

SHAP rankings were compared with age-associated gene sets from six independent studies: AFCA DEGs and clock genes [7], Gao et al. aging clock genes [8], Tabula Muris Senis DEGs and clock genes [6], and Zhu et al. PBMC variable-importance genes [19]. Enrichment is the observed overlap divided by the overlap expected under random selection within the shared gene set, with significance from a hypergeometric test.

### 4.10 Inference-time gene perturbation

Target genes were hidden from the test input while model weights stayed fixed. For scGPT the target genes were set to zero before tokenization, which drops them from the input sequence; for Geneformer their tokens were removed from the rank-ordered sequence. Six conditions were tested: the 25 RP genes in the SHAP top 50, all 51 RP genes, the top 50 SHAP genes, the top 100 SHAP genes, and size-matched sets of 25 or 51 randomly selected non-RP genes. Five random sets were drawn at each size, and *Z* scores were computed against the matching control distribution.

### 4.11 Differential expression analysis

Counts were aggregated by donor or sample and tested for pseudobulk differential expression with the robust quasi-likelihood method in edgeR. The Supplementary Information sets out the replicate structure for each species and marks which analyses support inference and which are descriptive.

## Supporting information

supplemental fies updated.

## Supplementary information

Supplementary figures, tables, and methods are available online.

## Acknowledgements

We used the Wahab HPC cluster at Old Dominion University for computation and Weights & Biases for experiment tracking.

## Declarations

### Ethics approval

Not applicable. This study used only publicly available, deidentified datasets.

### Data availability

Fly data from the Aging Fly Cell Atlas. Worm data from Gao et al. (GSE229022). Mouse data from Tabula Muris Senis (Figshare). Human PBMC from CZ CELLxGENE Discover.

### Code availability

Code will be made available upon publication.

### Author contributions

H.Q. conceived and designed the study, acquired funding, revised the code and analyses, and supervised the research. K.L. initiated the study, designed the data-preprocessing and scGPT-training pipelines, and drafted the first version of the manuscript. M.A.U.A.A. extended the scGPT analysis to the human dataset, incorporated Geneformer, performed the post-training analyses, and drafted the second version. All authors revised the manuscript.

