## supplemental fies updated. for "Single-cell foundation models identify shared and divergent transcriptomic signatures of aging across invertebrates and mammals"

**S5 Supplementary Tables** **11**

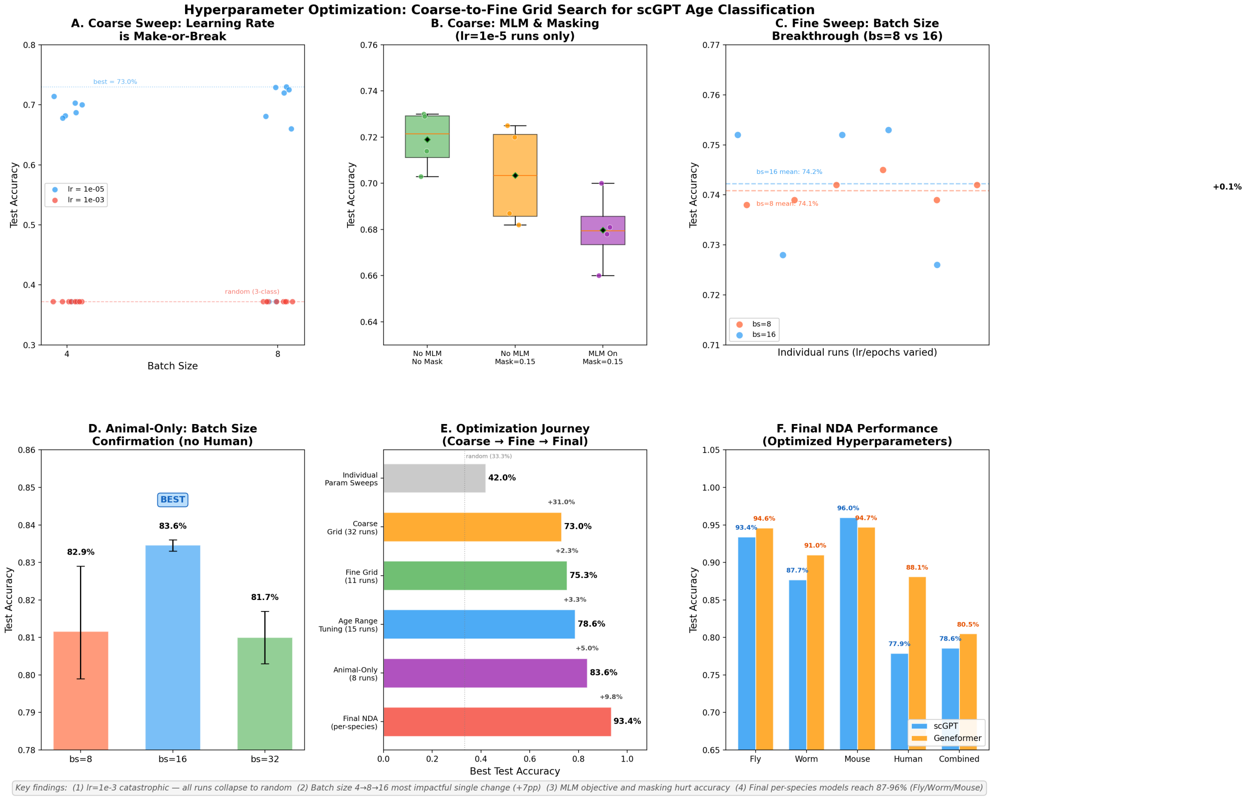

Figure S1: Four-stage coarse-to-fine hyperparameter search for scGPT. Across 66 runs, Combined NDA test accuracy increased from 42% to 78.6%. Learning rate and batch size had the largest effects on accuracy. The selected values were $1 \times 10^{-5}$ and 16, respectively; all runs with a learning rate of $1 \times 10^{-3}$ produced 37.2% accuracy. The masked-language-model objective and input masking were disabled during classification.

### S1 Donor-aware analysis

In addition to the non-donor-aware (NDA) models used for the primary SHAP analysis, we trained donor-aware (DA) models. All cells from a donor or sample were assigned to either the training set or the held-out test set. The DA models had lower test accuracy than the NDA models in every species (Table S1). This result shows that performance on cells from unseen donors was lower than performance under a cell-level split.

#### S1.1 Donor-aware differential expression

We performed donor- or sample-aware pseudobulk aggregation followed by robust quasi-likelihood testing in edgeR. The biological replicate was the donor for human, the individual mouse for mouse, and the sample for worm. For fly, we extracted sequencing-library identifiers from the cell barcodes. AFCA barcodes contain labels such as `female_body_70_S1`, whereas FCA barcodes contain source-library identifiers such as `FCA22_Female_body_adult`.

Table S1: Test accuracy under non-donor-aware (NDA) and donor-aware (DA) splits.

| | **scGPT** | | **Geneformer** | |
|---|---|---|---|---|
| **Species** | NDA | DA | NDA | DA |
| Fly | 93.4% | 68.6% | 94.6% | 51.4% |
| Worm | 87.7% | 61.0% | 91.0% | 65.1% |
| Mouse | 96.0% | 54.5% | 94.7% | 72.8% |
| Human | 77.9% | 66.1% | 88.1% | 51.4% |

Because these libraries represent independent pools of dissected flies, we used them as the replicate unit in the supplementary pseudobulk analysis.

Within each species, cells were stratified by annotated cell type. Raw counts were aggregated by summing across all cells belonging to the same donor, sample, or sequencing library and cell type, producing one pseudobulk count profile per biological replicate per cell type. Replicates contributing fewer than 20 cells to a given cell type were excluded. Strata with fewer than two biological replicates in either the young or old age group were excluded as underpowered.

For each eligible cell-type stratum, we constructed a `DGEList` from the pseudobulk count matrix, removed low-count genes with `filterByExpr`, and normalized library sizes by the trimmed mean of M-values method. We estimated dispersion with `estimateDisp`, fit the model with `glmQLFit` and `robust=TRUE`, and used `glmQLFTest` to compare old and young replicates.

#### S1.2 Donor-aware SHAP attribution

We used GradientExplainer to compute SHAP values for held-out cells from donors not represented in training. We sampled a fixed number of cells per donor and cell type, then aggregated cell-level SHAP values by donor within each cell type. For every gene and donor, we calculated the mean signed and mean absolute SHAP values across cells. We then calculated two summaries for each cell type:

$$\Delta\phi_g = \bar{\phi}_g^{\text{old}} - \bar{\phi}_g^{\text{young}}$$
$$\bar{A}_g = \tfrac{1}{2}\left(\bar{A}_g^{\text{old}} + \bar{A}_g^{\text{young}}\right)$$

where $\Delta\phi_g$ is the signed SHAP difference between old and young donors for gene $g$, and $\bar{A}_g$ is the mean absolute SHAP across all donors.

#### S1.3 Species-level applicability

The four datasets differed in donor structure and in the number of replicates retained after donor-aware splitting (Table S2). We therefore use the pseudobulk results as a supplementary comparison rather than as the primary validation of the SHAP rankings. Differential expression tests marginal changes in expression, whereas SHAP measures a gene's contribution to the model's age classification.

Table S2: Species-specific inferential versus descriptive status of downstream DEG and SHAP analyses.

| Species | Donor/sample unit | DEG mode | SHAP mode | Interpretation |
|---|---|---|---|---|
| Human | True donor ID (`donor_id`) | Donor-aware pseudobulk edgeR | Donor-level aggregation | Mostly inferential, but GIS-only restriction leaves limited old-donor support and few global DEGs. |
| Mouse | True mouse ID (`mouse.id`) | Donor-aware pseudobulk edgeR | Donor-level aggregation | Mixed; inferential only where mouse counts were adequate, with the 1m versus 30m contrast limited by two young mice. |
| Worm | True sample ID (`orig.ident/sample`) | Sample-aware pseudobulk edgeR | Sample-level aggregation where supported | DEG was inferential but minimally powered, with three young and three old biological replicates. |
| Fly | Barcode-derived sequencing library ID | Library-aware pseudobulk edgeR | Library-level or cell-type summary | Inferential as supplementary pseudobulk DEG using 26 young and 24 old libraries for 5d versus 70d. |

Because the donor-aware splits left few replicates for several age-by-species comparisons, the primary SHAP analysis uses the NDA models. The DA results provide a separate assessment of generalization to unseen donors and should be interpreted with their smaller sample sizes in mind.

### S2 Gene-removal ablation with retraining

The inference-time perturbation in the main text tests whether trained models rely on selected genes at prediction. Here, we removed genes *before* training and retrained both models from their pretrained checkpoints. This complementary analysis tests how well the models can use other features when the selected genes are absent throughout fine-tuning.

#### S2.1 Experimental design

We removed selected genes from the Combined NDA dataset (263,228 cells and 2,315 genes) and retrained both models with the same hyperparameters used for the corresponding baseline. We tested 17 conditions per model (34 runs in total): two RP-removal conditions, three top-$K$ SHAP conditions, and size-matched controls (Table S3; Fig. S2).

We identified RP genes with the pattern `^RPL\d+[A-Z]?$` | `^RPS\d+[A-Z]?$` | `^RPLP\d+$` | `^RPSA$`, excluding kinases such as RPS6KC1 and excluding RACK1. In each condition, we removed the same gene set from both models.

#### S2.2 Retraining ablation results

Table S3: Accuracy after gene removal and retraining. Delta is the change from baseline. Random controls report the mean ± SD across five independently sampled gene sets.

| **Condition (genes removed)** | **scGPT Acc** | Δ | **GF Acc** | Δ |
|---|---|---|---|---|
| Baseline (0) | 78.7% | — | 81.8% | — |
| *Top-K SHAP gene removal* | | | | |
| Top 10 (10) | 78.0% | −0.7 pp | 80.8% | −1.0 pp |
| Top 50 (50) | 77.4% | −1.3 pp | 79.7% | −2.1 pp |
| Top 100 (100) | 77.3% | −1.4 pp | 79.1% | −2.7 pp |
| *RP gene removal* | | | | |
| RP top 50 (25) | 77.3% | −1.4 pp | 78.5% | −3.3 pp |
| All RP (51) | 75.7% | −3.1 pp | 76.4% | −5.4 pp |
| *Controls* | | | | |
| Bottom 25 (25) | 78.2% | −0.5 pp | 81.1% | −0.7 pp |
| Random 25 mean (25) | 78.5% | −0.2 pp | 81.5% | −0.3 pp |
| Random 51 mean (51) | 78.7% | −0.0 pp | 81.3% | −0.5 pp |

Removing the 25 RP genes in the consensus top 50 reduced accuracy after retraining by 1.4 percentage points for scGPT and 3.3 points for Geneformer. Removing all 51 RP genes reduced accuracy by 3.1 and 5.4 points, respectively. The corresponding losses for randomly selected non-RP genes ranged from 0.0 to 0.5 points ($Z = 6.2$–$26.0$, $p < 0.001$).

The losses were much larger when the genes were hidden only at inference: 27.6 points for scGPT and 22.1 points for Geneformer. When the models were retrained without these genes, the losses were 3.1 and 5.4 points. This fourfold to ninefold difference indicates that the retrained models could make greater use of the remaining 2,264 non-RP genes. The inference-time experiment measures reliance by the original models; it does not establish a causal role for RP genes in aging.

Removing all 51 RP genes also reduced accuracy more than removing the top 100 SHAP genes: 3.1 versus 1.4 points for scGPT and 5.4 versus 2.7 points for Geneformer. Geneformer was more sensitive than scGPT to RP removal (5.4 versus 3.1 points), even though RP genes appeared less often in its invertebrate SHAP rankings. This difference suggests that the individual-gene rankings do not fully describe the information contributed by the RP set.

### S3 SHAP rankings after RP removal

We recalculated SHAP values after removing RP genes and retraining the models. This analysis shows how the rankings changed when RP genes were no longer available (Fig. S4;

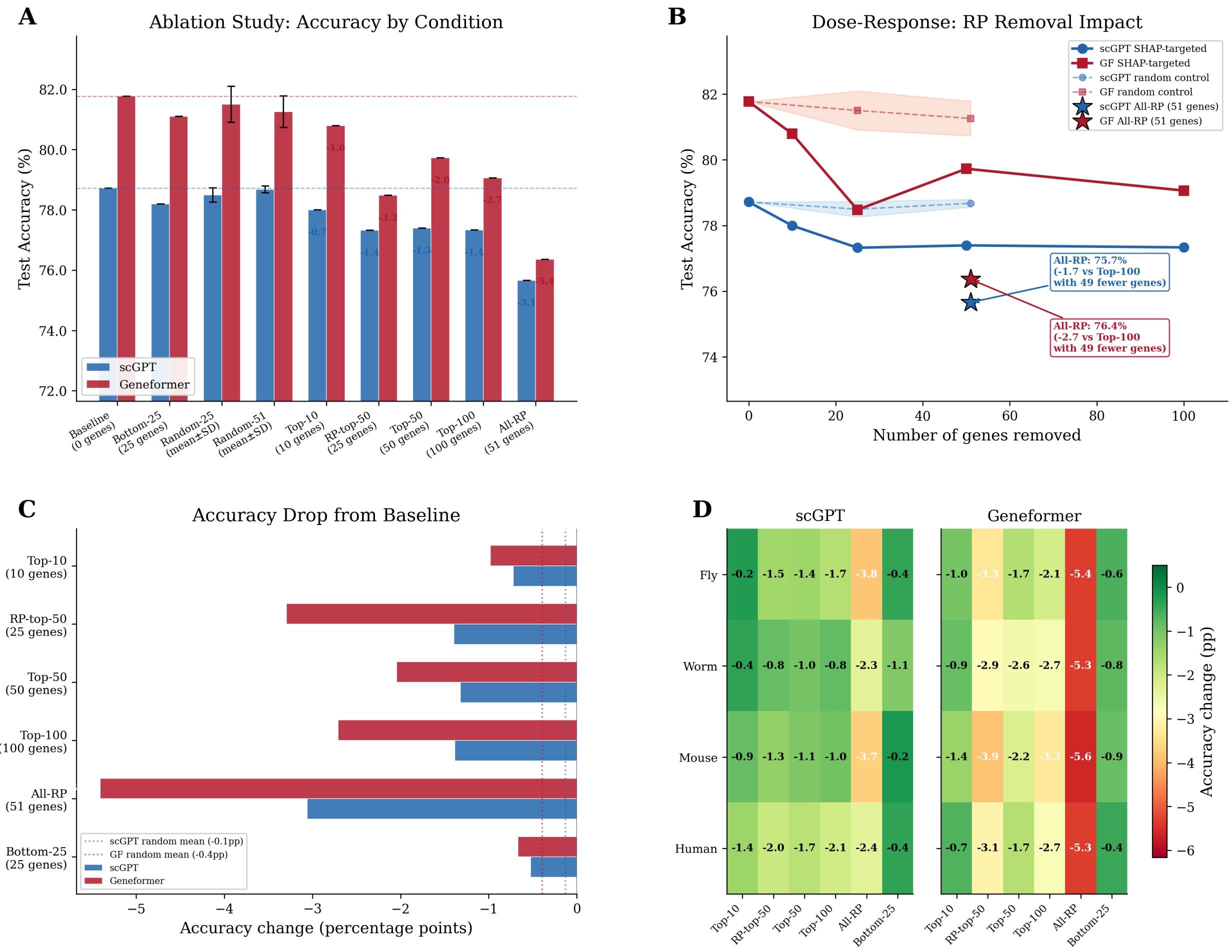

Figure S2: Accuracy after gene removal and retraining. (A) Accuracy by condition and model. (B) Accuracy as a function of the number of RP, top-$K$ SHAP, or random genes removed. Removing all 51 RP genes reduces accuracy more than removing the top 100 SHAP genes. (C) Change in accuracy relative to baseline. (D) Results by species and model.

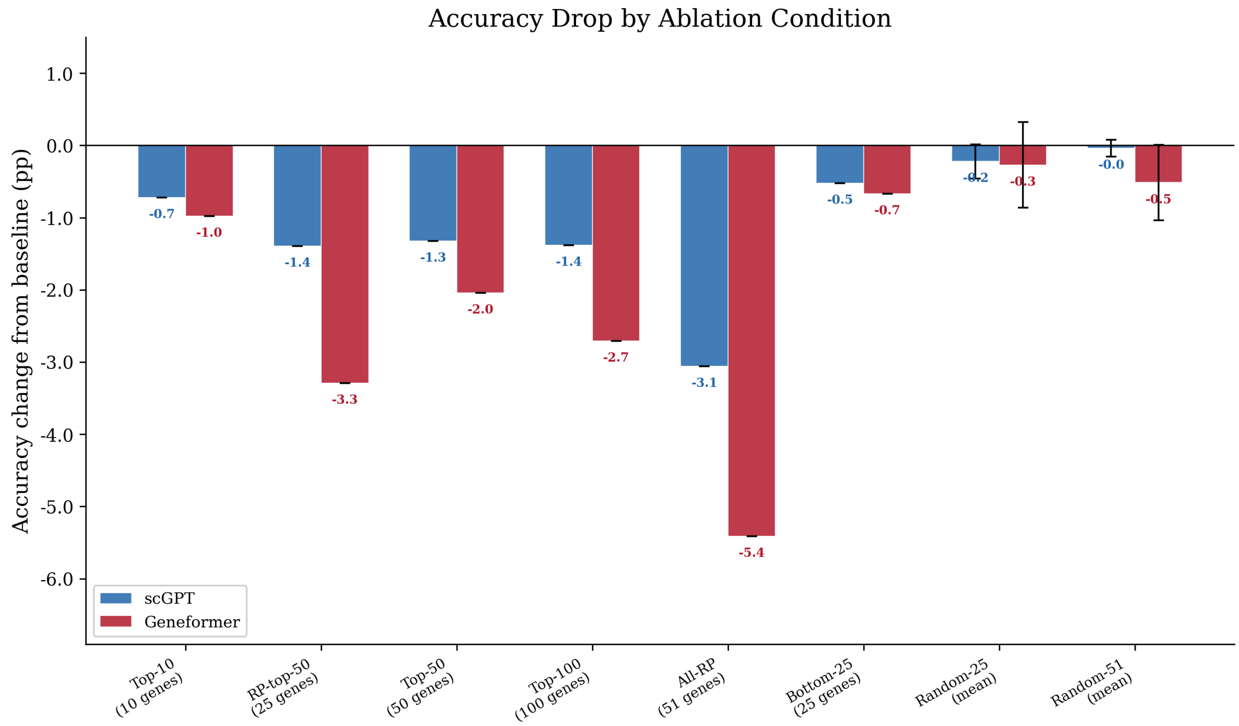

Figure S3: Change in accuracy after gene removal and retraining. Negative values indicate a loss relative to baseline (78.7% for scGPT and 81.8% for Geneformer). RP-gene removal produced larger losses than size-matched random controls. Error bars for random controls show ±1 SD across five independently sampled gene sets.

Table S4).

Table S4: Top 10 consensus genes after removing all 51 RP genes and retraining. Mean rank is calculated across the four species.

| Rank | scGPT gene | Mean rank | GF gene | Mean rank |
|---|---|---|---|---|
| 1 | OAZ1 | 1.00 | SPTBN1 | 12.75 |
| 2 | EEF2 | 3.75 | TCF25 | 20.75 |
| 3 | ARPC2 | 5.75 | CNOT4 | 33.00 |
| 4 | ARL6IP5 | 6.00 | CCT6A | 33.75 |
| 5 | HSP90B1 | 8.00 | TCF4 | 34.75 |
| 6 | POLG | 8.00 | SYNE1 | 40.00 |
| 7 | VAMP2 | 8.75 | NIPBL | 40.75 |
| 8 | ITPR1 | 11.25 | SPAG9 | 41.50 |
| 9 | STX16 | 12.00 | RALBP1 | 44.00 |
| 10 | PFDN5 | 13.00 | ELF2 | 50.00 |

**scGPT rankings after RP removal.** After we removed all 51 RP genes and retrained scGPT, OAZ1 (ornithine decarboxylase antizyme 1) ranked first in all four species. OAZ1 regulates polyamine metabolism upstream of mTOR and translational control, and the second-ranked gene, EEF2, functions in translational elongation. The prominence of these two genes suggests that translation-related features remained important after RP removal. Only 2 of the baseline top 50 non-RP genes remained in the post-ablation top 50, indicating that the scGPT ranking changed extensively.

**Geneformer rankings after RP removal.** Geneformer's post-ablation top 50 retained 32 genes from the baseline top 50. SPTBN1, which encodes a cytoskeletal scaffolding protein, ranked first in the consensus, and TCF25 remained second. The smaller change is consistent with the lower prevalence of RP genes in Geneformer's baseline invertebrate rankings.

**Partial RP removal.** When we removed only the 25 top-ranked RP genes, RPS27 became scGPT's highest-ranked gene in all four species. RPL27A, RPLP1, RPL38, and RPS20 also entered the top 10, showing that other RP genes carried similar predictive information. No RP gene appeared in Geneformer's top 10 under this condition; the highest-ranked genes instead included SYNE1, TCF25, RALBP1, TOMM7, and HECTD1.

### S4 Overlap between differential expression and SHAP rankings

We compared the SHAP rankings with differentially expressed genes (DEGs), but neither available DEG analysis provides a direct validation. A cell-level Wilcoxon test treats cells as independent and is vulnerable to pseudoreplication. Pseudobulk analysis in edgeR uses

**Post-Ablation SHAP: Gene Takeover After RP Removal**

**A** scGPT: Top-10 Consensus Genes

| Rank | Production | RP-top-50 removed | All-RP removed |
|---|---|---|---|
| 1 | RPL12 | RPS27 | OAZ1 |
| 2 | RPS21 | RPL27A | EEF2 |
| 3 | RPS13 | DYNC1H1 | ARPC2 |
| 4 | RPS26 | RPLP1 | ARL6IP5 |
| 5 | RPS15 | RPL38 | HSP90B1 |
| 6 | RPL3 | RPS20 | POLG |
| 7 | RPS3 | SEC62 | VAMP2 |
| 8 | RPL17 | PLCB4 | ITPR1 |
| 9 | TOMM7 | PFDN5 | STX16 |
| 10 | RPL28 | TCF25 | PFDN5 |

**B** Geneformer: Top-10 Consensus Genes

| Rank | Production | RP-top-50 removed | All-RP removed |
|---|---|---|---|
| 1 | TCF25 | SYNE1 | SPTBN1 |
| 2 | RPS13 | TCF25 | TCF25 |
| 3 | SPTBN1 | RALBP1 | CNOT4 |
| 4 | NIPBL | TOMM7 | CCT6A |
| 5 | HECTD1 | HECTD1 | TCF4 |
| 6 | SPAG9 | CEBPZ | SYNE1 |
| 7 | RALBP1 | PAFAH1B1 | NIPBL |
| 8 | RFC1 | SPAG9 | SPAG9 |
| 9 | CCT6A | CDC5L | RALBP1 |
| 10 | PAFAH1B1 | CNOT4 | ELF2 |

Ribosomal protein · Non-RP (in production top-10) · Newly promoted

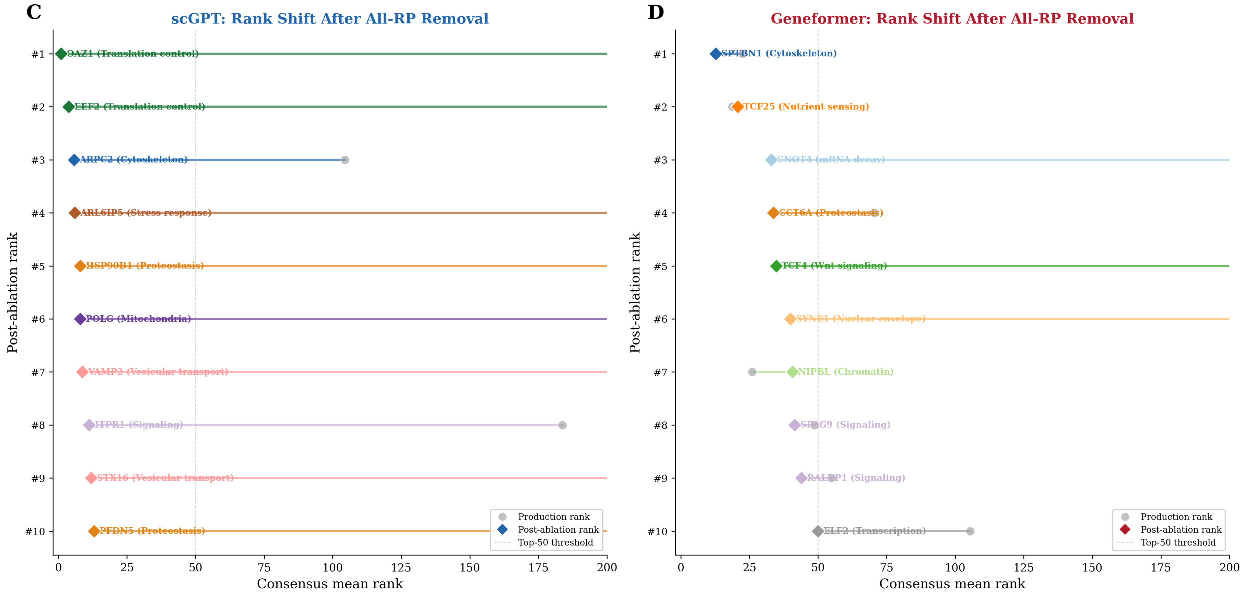

Figure S4: SHAP rankings after gene removal and retraining. (A, B) Top 10 consensus genes in the baseline, RP-top-50, and all-RP conditions for scGPT and Geneformer. Blue denotes RP genes, orange denotes non-RP genes present in the baseline top 10, and green denotes genes that enter the top 10 after removal. (C, D) Baseline ranks (gray circles) and post-ablation ranks (colored diamonds) for the top 10 genes after removal.

biological replicates but requires enough replicates in each age group. Under the NDA age definitions, the mouse data included only two 1-month-old mice, and the human data included 25 young and 147 old donors. We therefore changed the age boundaries for the pseudobulk comparison (Table S5). Because the pseudobulk and SHAP analyses use different age definitions for mouse and human, we present their overlap as a sensitivity analysis.

Table S5: Age-label adjustments for pseudobulk DEG analysis. Production NDA labels are those used to train the Combined model whose SHAP rankings are being validated.

| Species | Production NDA labels | Pseudobulk labels | Rationale |
|---|---|---|---|
| Fly | Young: 5d, Old: 70d | Young: 5d, Old: 70d | No change. 26 young and 4 old libraries. |
| Worm | Young: D1, Old: D14 | Young: D1, Old: D14 | No change. 3 young and 3 old replicates. |
| Mouse | Young: 1m, Old: 30m | Young: 3m, Old: 30m | Changed. Only 2 mice at 1m; 7 mice at 3m enable per-tissue replication. |
| Human | Young: 19–22y, Old: 50–75y | Young: 19–26y, Old: 60–75y | Changed. Ratio improved from 1:5.9 to 1.3:1 (78 vs 58 donors). |

#### S4.1 Pseudobulk methodology

We summed raw counts within each biological replicate (donor, mouse, or sample batch). For each species, we constructed a `DGEList`, filtered low-count genes with `filterByExpr`, normalized library sizes by the trimmed mean of M-values method, and estimated dispersions with empirical Bayes shrinkage. We used edgeR's exact test to compare old and young groups and controlled the false-discovery rate by the Benjamini–Hochberg procedure. We defined DEGs by FDR $< 0.05$ and $|\log_2 \mathrm{FC}| > 1.0$. For mouse, we analyzed the six tissues with at least two mice per group separately and took the union of their DEGs. Liver contributed 4,229 of the 4,779 mapped genes in this union.

#### S4.2 Pseudobulk DEG-SHAP overlap results

The top 50 SHAP genes were enriched among pseudobulk DEGs in seven of the eight species–model comparisons (Table S6), although the age definitions differed for mouse and human.

For fly, enrichment was 4.5–4.7-fold, and all genes overlapping the Geneformer ranking had the same direction of age effect in both analyses. Worm enrichment was 2.4–2.9-fold, with 82–100% directional agreement, but each age group had only three replicates. Mouse enrichment was 2.0–2.1-fold and directional agreement was 17–36%. The low agreement may reflect the union of genes with tissue-specific effects. Human scGPT had no overlap

Table S6: Pseudobulk DEG-SHAP overlap at $K = 50$ (Combined NDA model, $|\log_2 \mathrm{FC}| > 1.0$, FDR $< 0.05$). Enrichment is fold over hypergeometric baseline. Dir. % indicates the fraction of overlapping genes with concordant SHAP and DEG age-effect direction.

| **Species** | **Model** | **Overlap** | **%** | **Enrich.** | ***p*-value** | **Dir. %** |
|---|---|---|---|---|---|---|
| Fly | scGPT | 28/50 | 56.0 | 4.7× | $3.4 \times 10^{-14}$ | 68 |
| Fly | Geneformer | 27/50 | 54.0 | 4.5× | $3.4 \times 10^{-13}$ | 100 |
| Worm | scGPT | 11/50 | 22.0 | 2.4× | $5.1 \times 10^{-3}$ | 82 |
| Worm | Geneformer | 14/50 | 28.0 | 2.9× | $1.4 \times 10^{-4}$ | 100 |
| Mouse | scGPT | 29/50 | 58.0 | 2.1× | $5.9 \times 10^{-6}$ | 17 |
| Mouse | Geneformer | 28/50 | 56.0 | 2.0× | $2.3 \times 10^{-5}$ | 36 |
| Human | scGPT | 0/50 | 0.0 | 0.0× | 1.0 | — |
| Human | Geneformer | 4/50 | 8.0 | 13.8× | $1.4 \times 10^{-4}$ | 75 |

with the 206 DEGs. Four human Geneformer genes overlapped, corresponding to 13.8-fold enrichment.

The seven significant comparisons had enrichments of 2.0–13.8-fold ($p < 0.05$). The changed age labels for mouse and human limit direct comparison with the SHAP analysis, so these results are supplementary to the main validation analyses.

### S5 Supplementary Tables

Table S7: Summary of age-relabeled fine-tuning settings and test accuracy for *scGPT* and *Geneformer* across single species, pairwise, and pooled training regimes. Age classes were relabeled as `young`, `middle`, and `old`. For the human-only model (Model 4), donor-aware accuracy is reported; all other models report test accuracy under a cell-level split (see Methods).

| | *Model 1* | *Model 2* | *Model 3* | *Model 4* (Donor-aware) | *Model 5* | *Model 6* | *Model 7* | *Model 8* | *Model 9* | *Model 10* | *Model 11* | *Model 12* |
|---|---|---|---|---|---|---|---|---|---|---|---|---|
| Training setting | Fly only | Worm only | Mouse only | Human only | Fly + Worm | Worm + Mouse | Fly + Mouse | Fly + Human | Worm + Human | Mouse + Human | Pooled (Fly+Worm +Mouse) | Pooled (Fly+Worm +Mouse+Human) |
| Fly | ✓ | | | | ✓ | | ✓ | ✓ | | | ✓ | ✓ |
| Worm | | ✓ | | | ✓ | ✓ | | | ✓ | | ✓ | ✓ |
| Mouse | | | ✓ | | | ✓ | ✓ | | | ✓ | ✓ | ✓ |
| Human | | | | ✓ | | | | ✓ | ✓ | ✓ | | ✓ |
| Classes to predict (Age relabeled) | `young`: 5d `middle`: 30d `old`: 70d | `young`: D1 `middle`: D6 `old`: D14 | `young`: 1m `middle`: 18m `old`: 30m | `young`: 19–22 `middle`: 38–42 `old`: 50–75 | `young` `middle` `old` | `young` `middle` `old` | `young` `middle` `old` | `young` `middle` `old` | `young` `middle` `old` | `young` `middle` `old` | `young` `middle` `old` | `young` `middle` `old` |
| scGPT Max. seq. length | 4417 | 3569 | 13939 | 6626 | 2421 | 3357 | 4246 | 4534 | 3569 | 13946 | 2338 | 2338 |
| **scGPT accuracy (%)** | 93.4 | 87.7 | 96.0 | 66.1 | 85.5 | 83.5 | 88.2 | 81.3 | 76.8 | 83.5 | 83.5 | 78.6 |
| **Geneformer accuracy (%)** | 94.6 | 91.0 | 94.7 | 51.4 | 85.8 | 87.8 | 89.8 | 83.9 | 78.3 | 87.4 | 84.2 | 80.5 |

Table S8: Human age labeling strategies and evaluation metrics.

| | *Model 1* | *Model 2* | *Model 3* |
|---|---|---|---|
| Human age setting | Every age class included | Relabeled (fixed ranges) | Single age per class |
| Age class definition | All observed ages (no relabeling) | Young: (19.0 – 22.0)<br>Middle: (38.0 – 42.0)<br>Old: (50.0 – 75.0) | Young: 21.0<br>Middle: 43.0<br>Old: 61.0 |
| Accuracy | 17.8 | 77.90 | 79.10 |
| Config (Batch Size) | 4 | 16 | 16 |

Table S9: Cross-species prediction performance. Each model was trained on one species and evaluated on the other three.

| | *Model 1* | *Model 2* | *Model 3* | *Model 4* |
|---|---|---|---|---|
| Trained on | Fly | Worm | Mouse | Human |
| Overall accuracy | 0.294 | 0.413 | 0.332 | 0.299 |
| Overall precision | 0.301 | 0.411 | 0.314 | 0.316 |
| Overall recall | 0.310 | 0.420 | 0.323 | 0.301 |
| Overall macro F1 | 0.237 | 0.394 | 0.211 | 0.290 |

Table S10: Summary of scGPT fine-tuning settings and performance by species, with age relabeled as `young`, `middle`, and `old`.

| | *Model 1* | *Model 2* | *Model 3* | *Model 4* | *Model 5* | *Model 6* | *Model 7* | *Model 8* | *Model 9* | *Model 10* | *Model 11* | *Model 12* |
|---|---|---|---|---|---|---|---|---|---|---|---|---|
| Training setting | Fly only | Worm only | Mouse only | Human only | Fly + Worm | Worm + Mouse | Fly + Mouse | Fly + Human | Worm + Human | Mouse + Human | Pooled (Fly+Worm +Mouse) | Pooled (Fly+Worm +Mouse+Human) |
| Fly | $\chi$ | | | | $\chi$ | | $\chi$ | $\chi$ | | | $\chi$ | $\chi$ |
| Worm | | $\chi$ | | | $\chi$ | $\chi$ | | | $\chi$ | | $\chi$ | $\chi$ |
| Mouse | | | $\chi$ | | | $\chi$ | $\chi$ | | | $\chi$ | $\chi$ | $\chi$ |
| Human | | | | $\chi$ | | | | $\chi$ | $\chi$ | $\chi$ | | $\chi$ |
| Classes to predict (Age relabeled) | `young`: 5d<br>`middle`: 30d<br>`old`: 70d | `young`: D1<br>`middle`: D6<br>`old`: D14 | `young`: 1m<br>`middle`: 18m<br>`old`: 30m | Young: (19.0 – 22.0)<br>Middle: (38.0 – 42.0)<br>Old: (50.0 – 75.0) | `young`<br>`middle`<br>`old` | `young`<br>`middle`<br>`old` | `young`<br>`middle`<br>`old` | `young`<br>`middle`<br>`old` | `young`<br>`middle`<br>`old` | `young`<br>`middle`<br>`old` | `young`<br>`middle`<br>`old` | `young`<br>`middle`<br>`old` |
| Max. Seq. Length | 4417 | 3569 | 13939 | 6626 | 2421 | 3357 | 4246 | 4534 | 3569 | 13946 | 2338 | 2338 |
| Accuracy | 93.4 | 87.7 | 96 | 77.9 | 85.5 | 83.5 | 88.2 | 81.3 | 76.8 | 83.5 | 83.5 | 78.6 |
| Accuracy (Donor_Aware) | 68.6 | 61.0 | 54.5 | 66.1 | 71.7 | 73.1 | 67.2 | 64.6 | 72.2 | 78.6 | 68.7 | 67.21 |

Table S11: Summary of Geneformer donor-aware fine-tuning settings and performance by species, with age relabeled as `young`, `middle`, and `old`.

| | *Model 1* | *Model 2* | *Model 3* | *Model 4* | *Model 5* | *Model 6* | *Model 7* | *Model 8* | *Model 9* | *Model 10* | *Model 11* | *Model 12* |
|---|---|---|---|---|---|---|---|---|---|---|---|---|
| Training setting | Fly only | Worm only | Mouse only | Human only | Fly + Worm | Worm + Mouse | Fly + Mouse | Fly + Human | Worm + Human | Mouse + Human | Pooled (Fly+Worm +Mouse) | Pooled (Fly+Worm +Mouse+Human) |
| Fly | $\chi$ | | | | $\chi$ | | $\chi$ | $\chi$ | | | $\chi$ | $\chi$ |
| Worm | | $\chi$ | | | $\chi$ | $\chi$ | | | $\chi$ | | $\chi$ | $\chi$ |
| Mouse | | | $\chi$ | | | $\chi$ | $\chi$ | | | $\chi$ | $\chi$ | $\chi$ |
| Human | | | | $\chi$ | | | | $\chi$ | $\chi$ | $\chi$ | | $\chi$ |
| Classes to predict (Age relabeled) | `young`: 5d `middle`: 30d `old`: 70d | `young`: D1 `middle`: D6 `old`: D14 | `young`: 1m `middle`: 18m `old`: 30m | Young: (19.0 – 22.0) Middle: (38.0 – 42.0) Old: (50.0 – 75.0) | `young` `middle` `old` | `young` `middle` `old` | `young` `middle` `old` | `young` `middle` `old` | `young` `middle` `old` | `young` `middle` `old` | `young` `middle` `old` | `young` `middle` `old` |
| Accuracy | 94.6 | 91.0 | 94.7 | 88.1 | 85.8 | 87.8 | 89.8 | 83.9 | 78.3 | 87.4 | 84.2 | 80.5 |
| Accuracy (Donor_Aware) | 51.4 | 65.1 | 72.8 | 51.4 | 61.2 | 63.5 | 58.2 | 55.2 | 65.5 | 68.4 | 59.0 | 59.9 |
